## Supplementary material for "Resonating neurons stabilize heterogeneous grid-cell networks": 13 Figure supplements and 1 table supplement.

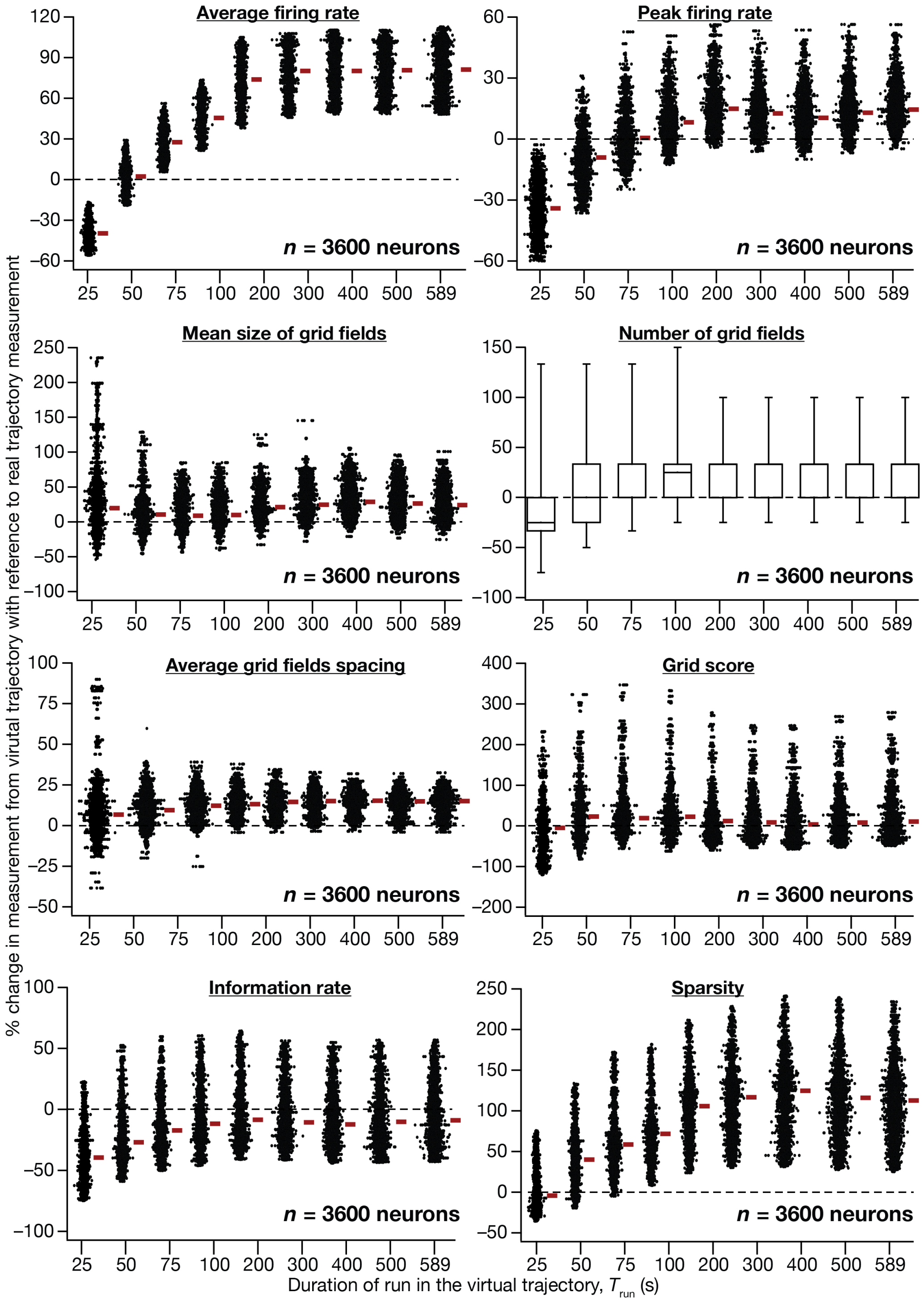

**Figure 1–figure supplement 1. Quantitative comparison of grid-cell firing properties obtained from CAN models simulated with virtual *vs.* real trajectories.** Percentage changes in average firing rate, peak firing rate, mean size, number, average grid field spacing, grid score, information rate, sparsity of grid-field rate maps obtained with a virtual trajectory for the 9 different values of $T_{run}$, in comparison to grid field rate maps obtained with the real trajectory ($T_{run}$=589 s). Values are shown for all neurons ($n$=3600) in the CAN model, and comparisons are made across respective neurons employing real *vs.* virtual trajectories.

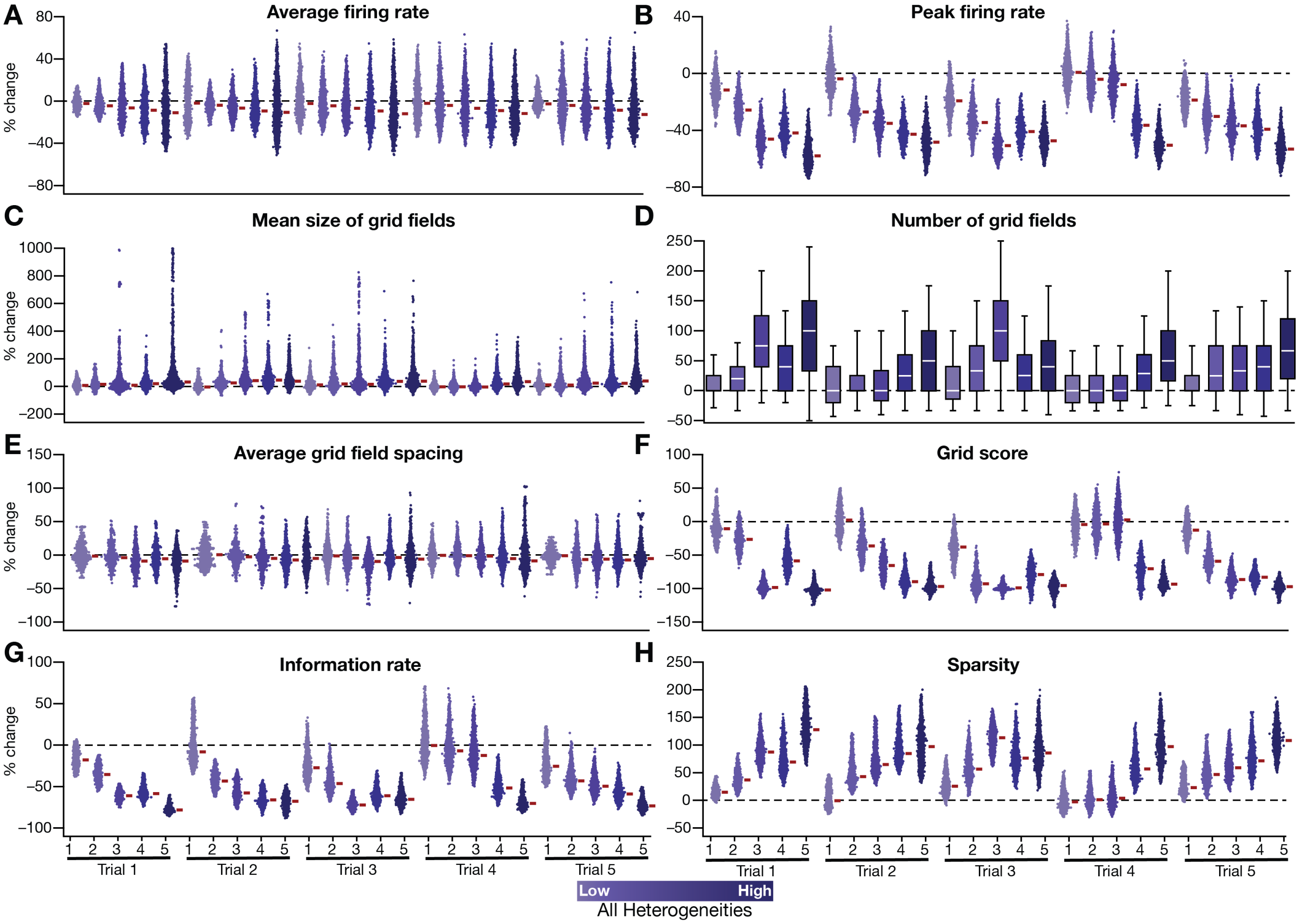

**Figure 3–figure supplement 1. Quantification of the disruption of grid-cell firing by network heterogeneities across different trials of CAN-model simulations.** (**A**–**H**) Grid cell activity of individual neurons in the network was quantified by 8 different measurements from five sets of trials (excluding the trial shown and quantified in Figure. 2-3). Distinct trials are obtained by unique initialization of the continuous attractor network and independent randomization of parametric values in introducing network heterogeneities. Depicted are percentage changes in each measurement of individual neurons (*n*=3600) in networks endowed with five degrees of heterogeneities, compared to the grid maps of respective neurons in the homogeneous network. All simulations in this figure are endowed with all the three forms (intrinsic, afferent and synaptic) of heterogeneities.

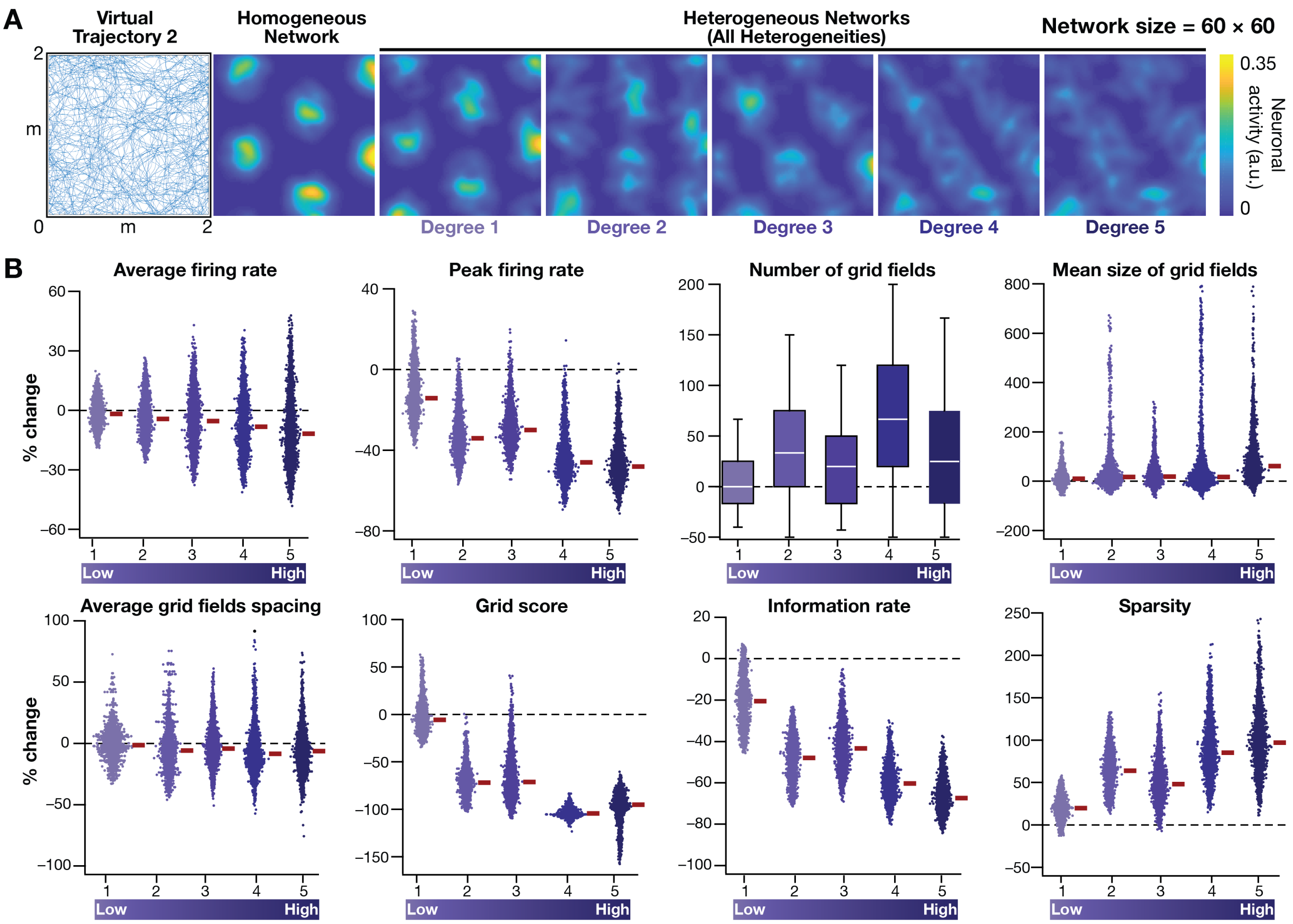

**Figure 3–figure supplement 2. Disruption of grid-cell firing by network heterogeneities was invariant to the specific trajectory employed by the CAN models.** (**A**) *Left to right*: Virtual trajectory in a 2 m×2 m square arena, which is distinct from that shown in Figure. 2*E*, was employed to perform CAN model simulations reported in this figure. Example rate map of grid cell activity in a homogeneous network, and in heterogeneous networks endowed with 5 degrees of heterogeneities, obtained with "Virtual trajectory 2". (**B**) Percentage changes in grid-cell measurements from heterogeneous CAN models with reference to those in homogeneous CAN model measurements for all neurons in the network (*n*=3600), obtained with "Virtual trajectory 2". All simulations in this figure are endowed with all the three forms (intrinsic, afferent and synaptic) of heterogeneities.

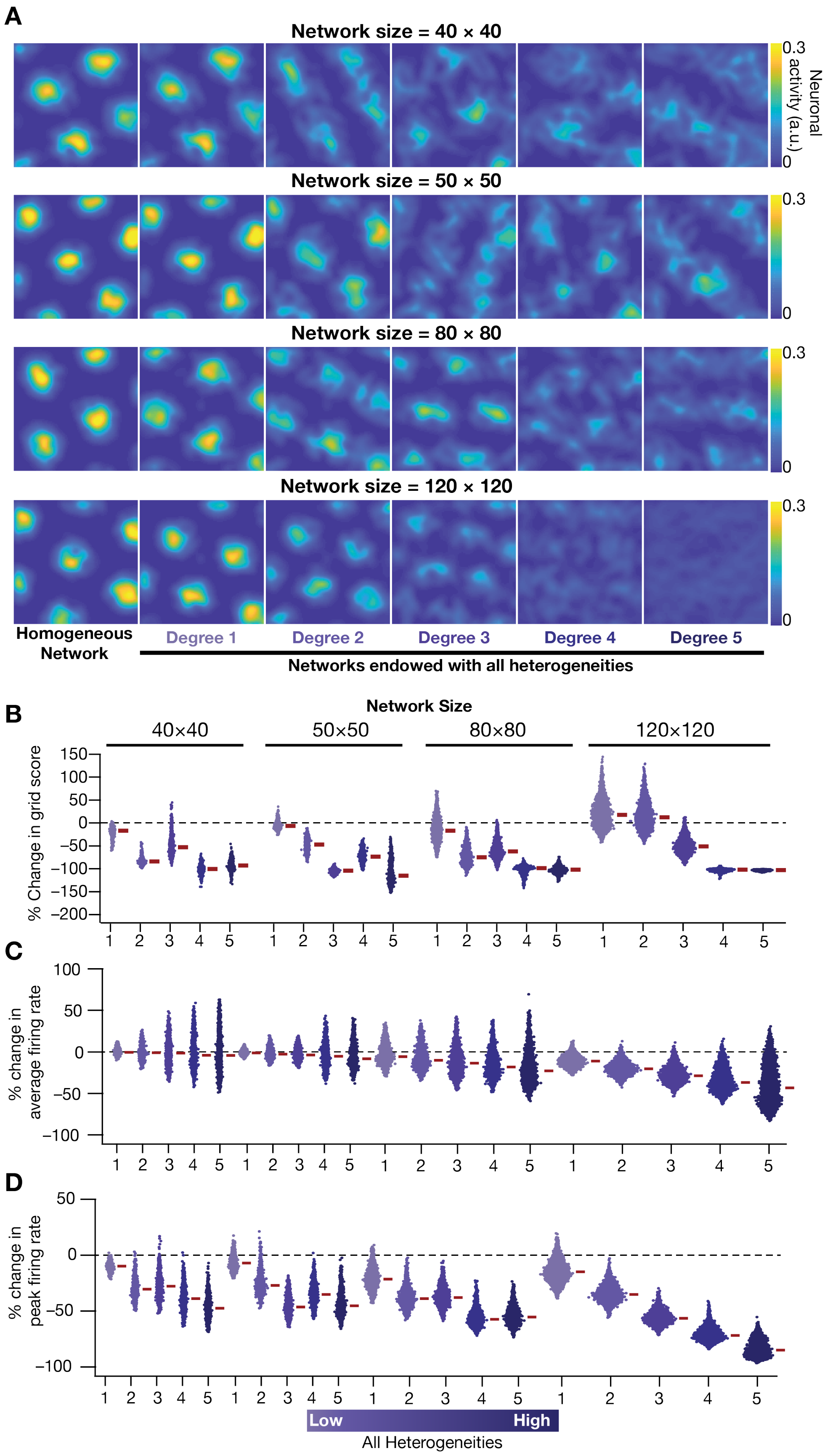
A
Network size = 40 × 40
Network size = 50 × 50
Network size = 80 × 80
Network size = 120 × 120
0.3
0
Neuronal activity (a.u.)
Homogeneous Network
Degree 1
Degree 2
Degree 3
Degree 4
Degree 5
Networks endowed with all heterogeneities
B
Network Size
40×40
50×50
80×80
120×120
% Change in grid score
C
% change in average firing rate
D
% change in peak firing rate
Low
High
All Heterogeneities

**Figure 3–figure supplement 3. Disruption of grid-cell activity by network heterogeneities was prevalent across CAN models of different sizes.** (**A**) Example rate maps of grid cell activity for homogeneous (*Column 1*), and heterogeneous networks endowed with 5 different degrees of heterogeneities (*Columns 2–6*) for networks of different sizes (*Row 1*: 40×40; *Row 2*: 50×50; *Row 3*: 80×80; *Row 4*: 120×120). (**B**) Percentage change in the grid score of individual neurons in networks of different sizes, endowed with five degrees of heterogeneities, compared to the grid score of neurons in homogeneous networks of respective sizes. Percent changes in average (**C**) and peak (**D**) firing rate of grid cell activity for individual neurons in heterogeneous network with different degrees of heterogeneity, computed with reference to the values from their respective homogeneous network. All simulations in this figure are endowed with all the three forms (intrinsic, afferent and synaptic) of heterogeneities.

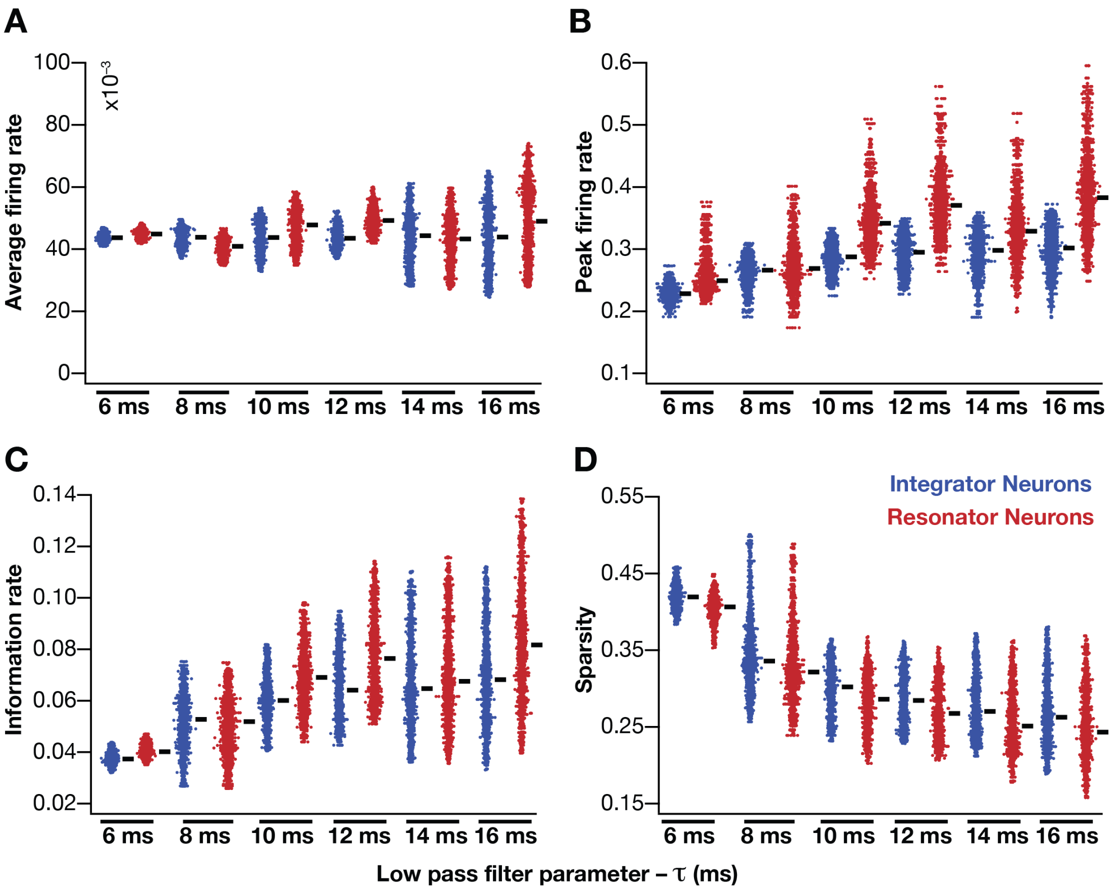

**Figure 6–figure supplement 1. Impact of neuronal resonance (phenomenological model), introduced by altering low-pass filter characteristics, on grid-cell characteristics in a homogeneous CAN model.** (**A**–**D**) average firing rate (**A**), peak firing rate (**B**), information rate (**C**) and sparsity (**D**) of grid fields in the arena for all neurons ($n$=3600) in homogeneous CAN models with integrator (blue) or resonator (red) neurons, modeled with different $\tau$ values. The HPF exponent $\varepsilon$ was set to 0.3 for all resonator neuronal models.

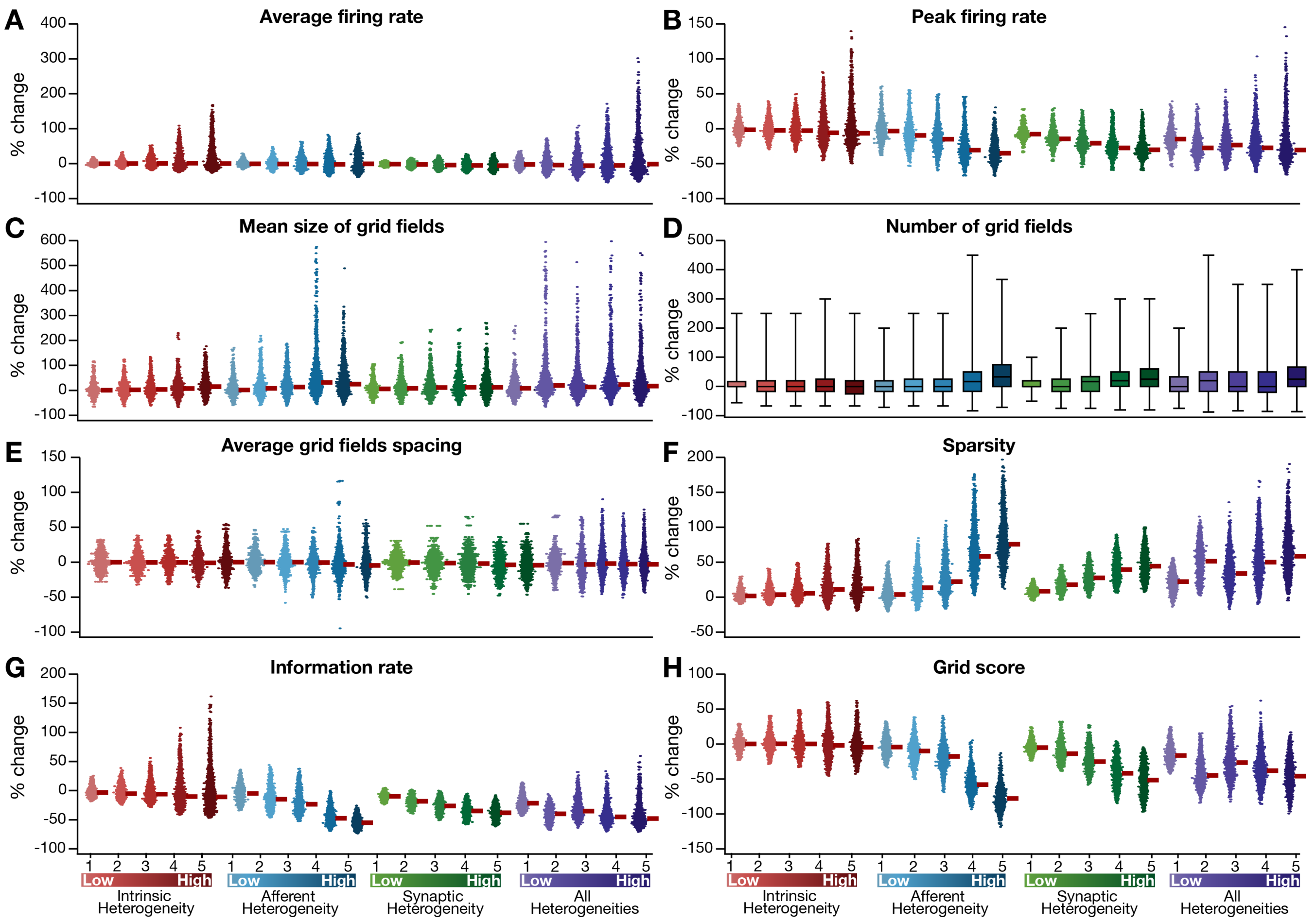

**Figure 8–figure supplement 1. Quantification of the grid-cell activity in presence of different forms of network heterogeneities in the CAN model with phenomenological resonator neurons.** Grid cell activity of individual resonator neurons in the network was quantified by 8 different measurements, for CAN models endowed independently with intrinsic, afferent or synaptic heterogeneities or a combination of all three heterogeneities. (**A**–**H**) Depicted are percentage changes in each of average firing rate (**A**), peak firing rate (**B**), mean size (**C**), number (**D**), average spacing (**E**), information rate (**F**), sparsity (**G**) and grid score (**H**) for individual neurons ($n$=3600) in networks endowed with distinct forms of heterogeneities, compared to the grid score of respective neurons in the homogeneous network.

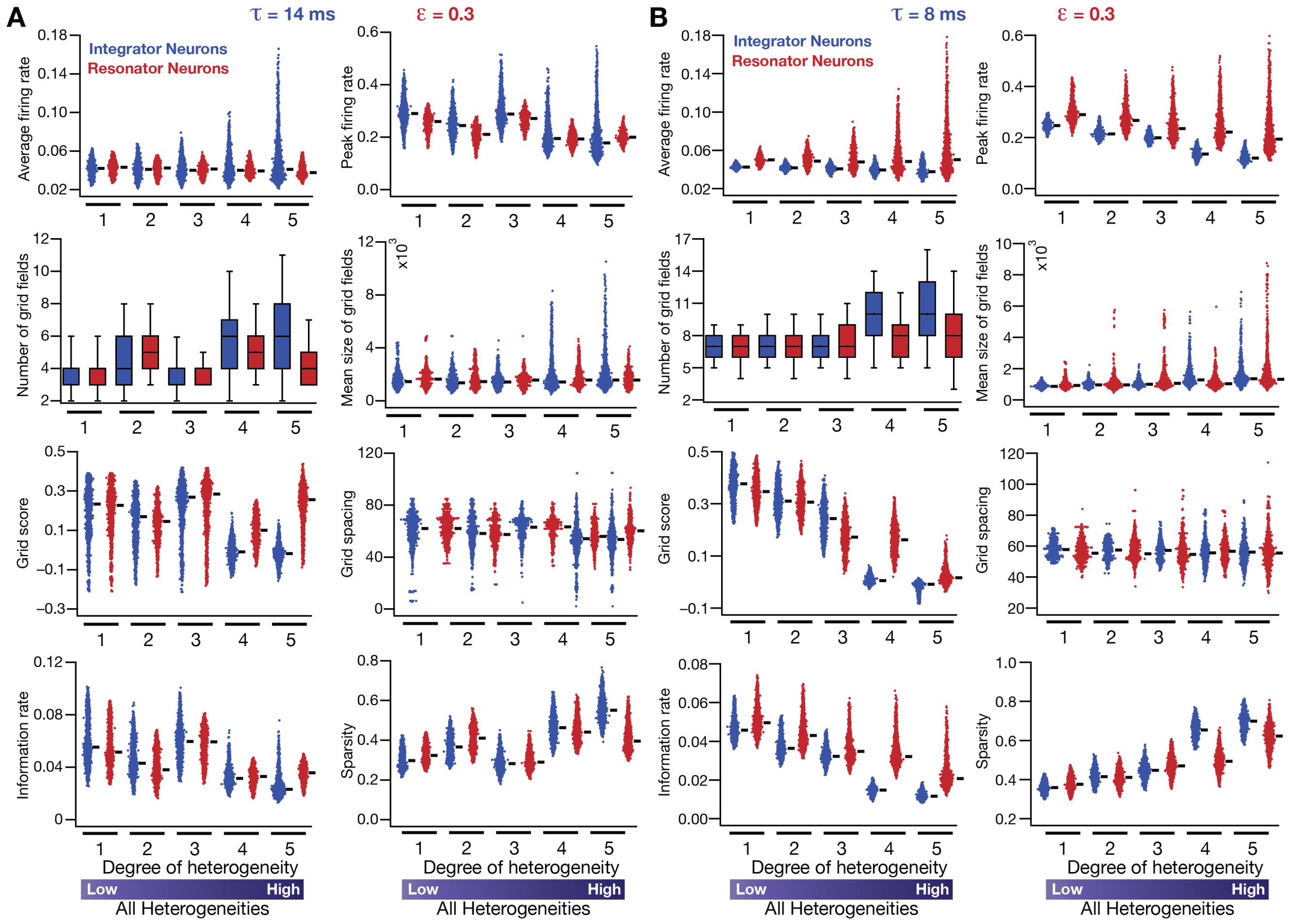

**Figure 8–figure supplement 2. Neuronal resonance (phenomenological) stabilizes grid-cell firing in heterogeneous CAN models.** (**A**) Average firing rate, peak firing rate, mean size, number, average grid field spacing, grid score, information rate and sparsity of grid fields for all neurons ($n$=3600) in heterogeneous CAN models with integrator (blue) or resonator (red) neurons, shown across 5 degrees of heterogeneities. All neurons in all networks were endowed with an integration time constant $\tau$ = 14 ms, and all resonator neurons were built with HPF exponent value $\varepsilon$ = 0.3. (**B**) Same as (**A**), but with $\tau$ = 8 ms for all neurons. All simulations in this figure are endowed with all the three forms (intrinsic, afferent and synaptic) of heterogeneities. In imposing intrinsic heterogeneities, the span of the uniform distributions that governed $\tau$ was always centered at the respective values ($\tau$ = 14 ms for $A$ and $\tau$ = 8 ms for $B$), with the extent of the uniform distribution increasing with increased degree of heterogeneity.

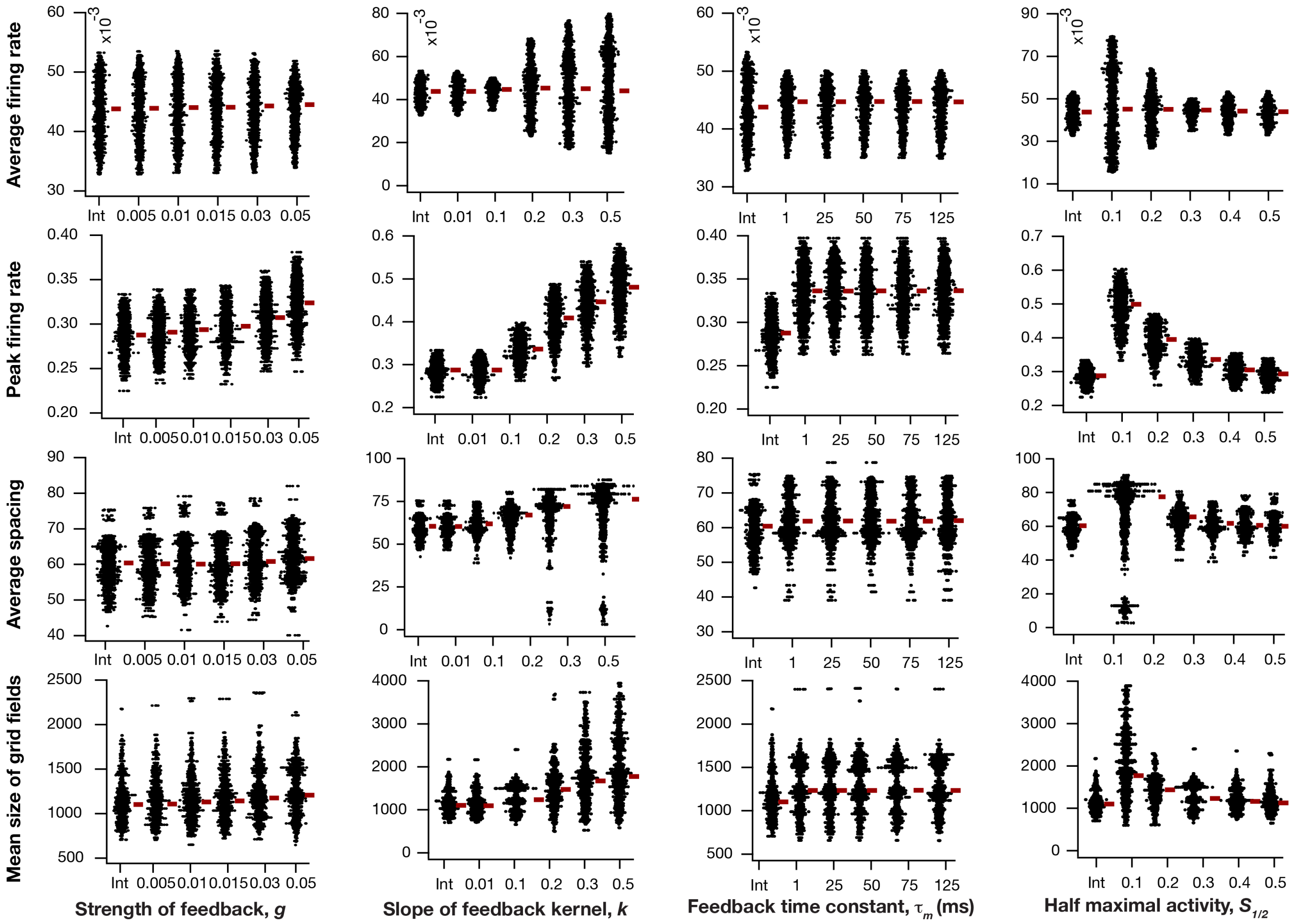

**Figure 10–figure supplement 1. Impact of intrinsic neuronal resonance, introduced by adding a negative feedback loop in the neuronal dynamics, on grid-cell characteristics in a homogeneous CAN model.** Average firing rate (*Row 1*), peak firing rate (*Row 2*), average spacing (*Row 3*) and mean size (*Row 4*) of grid fields in the arena for all neurons (*n*=3600) in homogeneous CAN models and their dependence on the parameters of negative feedback loop ($S_{1/2}$, $k$, $\tau_m$ and $g$) compared to homogeneous CAN model with integrator neurons (Int). Please note that the parameters were adjusted such that average firing rate in maintained at similar level across different parametric values.

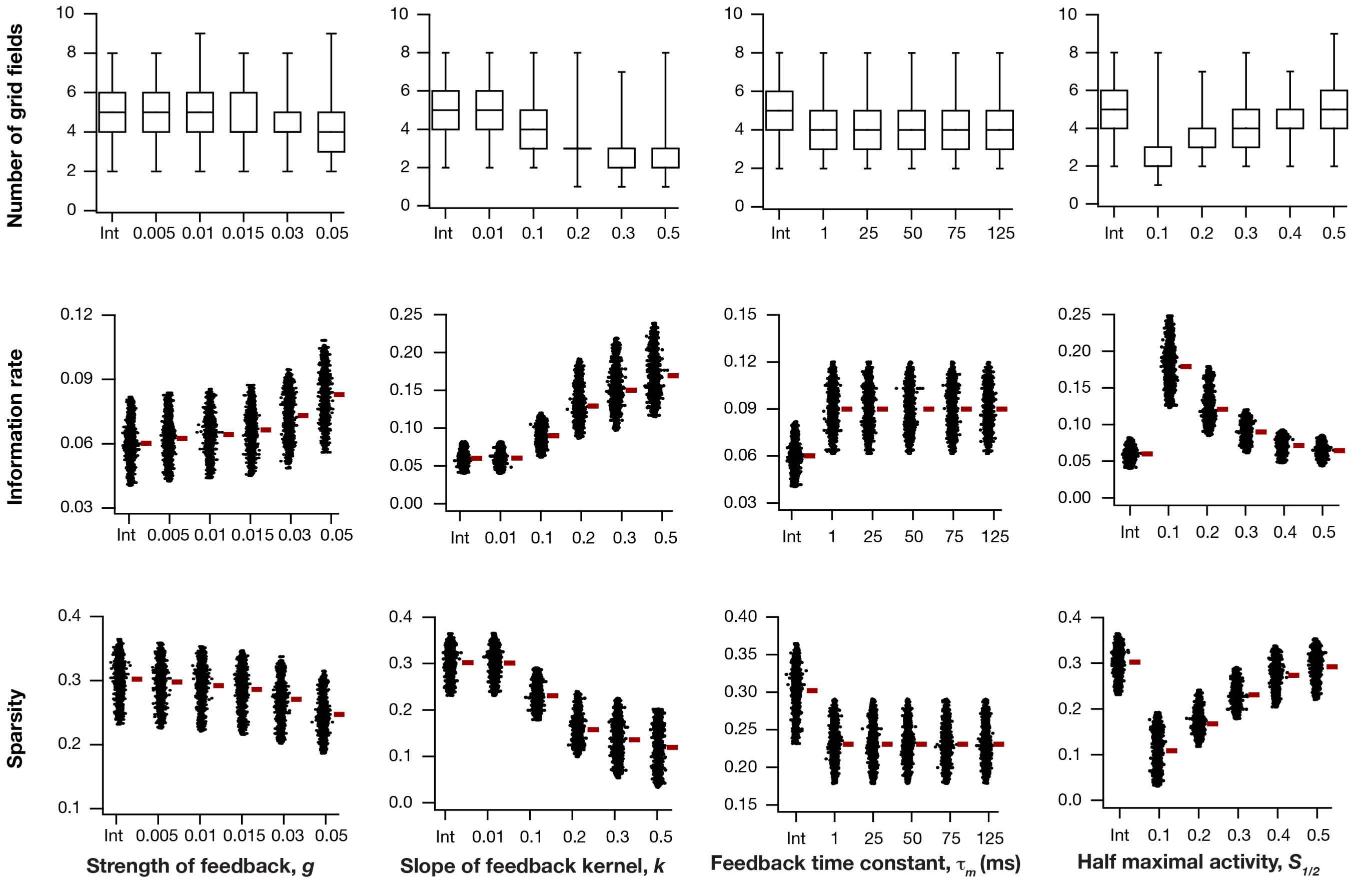

**Figure 10–figure supplement 2. Impact of intrinsic neuronal resonance, introduced by adding a negative feedback loop in the neuronal dynamics, on grid-cell characteristics in a homogeneous CAN model.** Number (*Row 1*), information rate (*Row 2*) and sparsity (*Row 3*) of grid fields in the arena for all neurons ($n$=3600) in homogeneous CAN models and their dependence on the parameters of negative feedback loop ($S_{1/2}$, $k$, $\tau_m$ and $g$) compared to homogeneous CAN model with integrator neurons (Int).

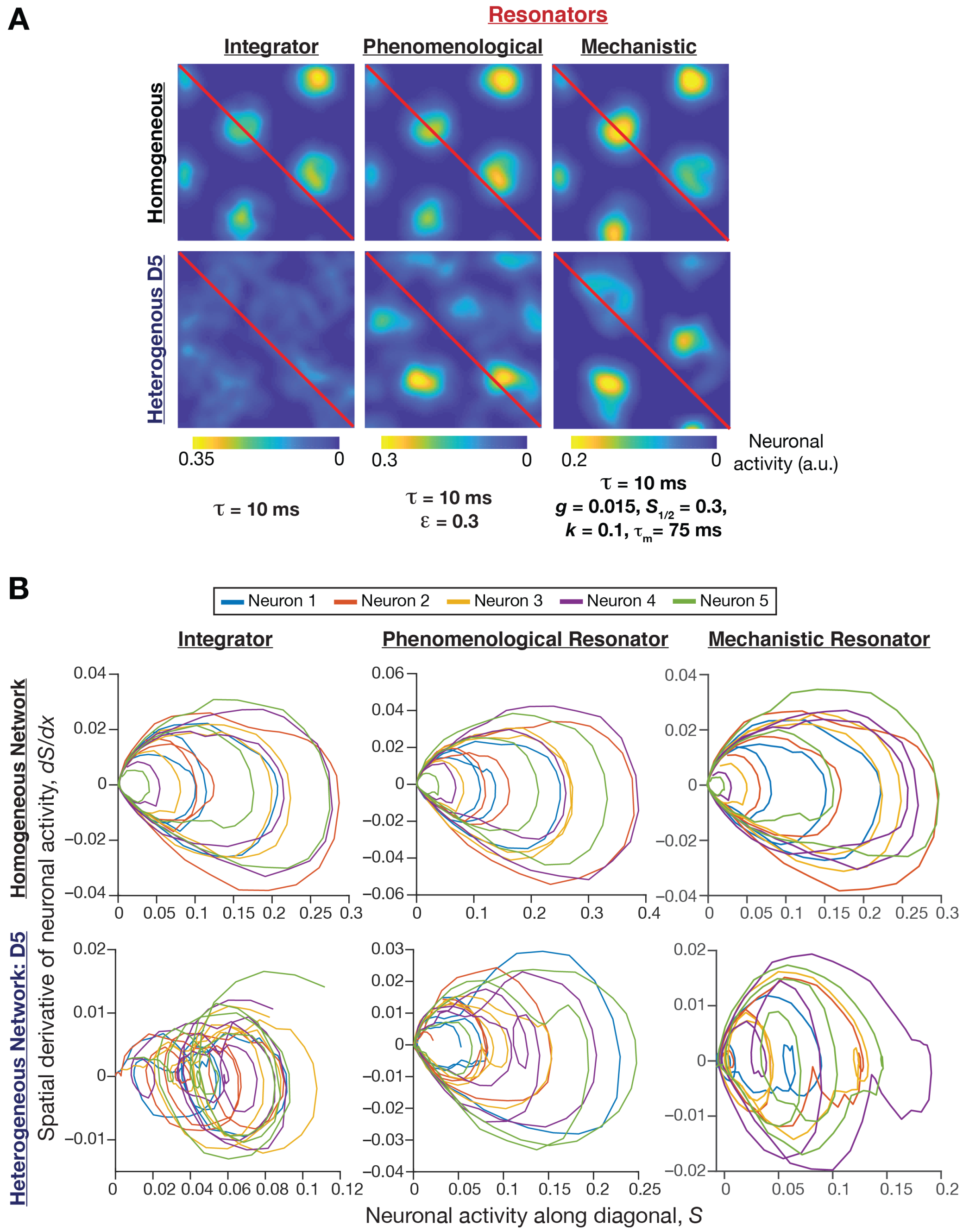

**Figure 11–figure supplement 1. Phase plane analysis of spatial profiles provided visualizations of the disruption of grid-cell activity in heterogeneous integrator networks and the relative robustness of heterogeneous resonator networks.** (**A**) Example rate maps of grid-cell activity in homogeneous (*Row 1*) and heterogeneous (*Row 2*; all heterogeneities, degree 5) CAN models, endowed with integrator (*Column 1*) or phenomenological resonator (*Column 2*) or mechanistic resonator (*Column 3*) neurons. Maps in *Columns 1–3* are from Figure 2*E*, Figure 8*A*, and Figure 11*A*, respectively. The diagonal values of these spatial rate maps (red lines) were employed for phase plane analysis. The choice of the diagonal was driven by the need to maximize the number of spatial activity cycles. (**B**) Phase-plane plots showing neuronal activity along the diagonal plotted against its spatial derivative for 5 different neurons in homogeneous (*Row 1*) and heterogeneous (*Row 2*; all heterogeneities, degree 5) CAN models, endowed with integrator (*Column 1*) or phenomenological resonator (*Row 2*) or mechanistic resonator (*Row 3*) neurons. Note the manifestation of closed orbits in homogeneous networks, the complete loss of closed orbits in the heterogeneous integrator network and the presence of noisy closed orbits in the heterogeneous resonator networks.

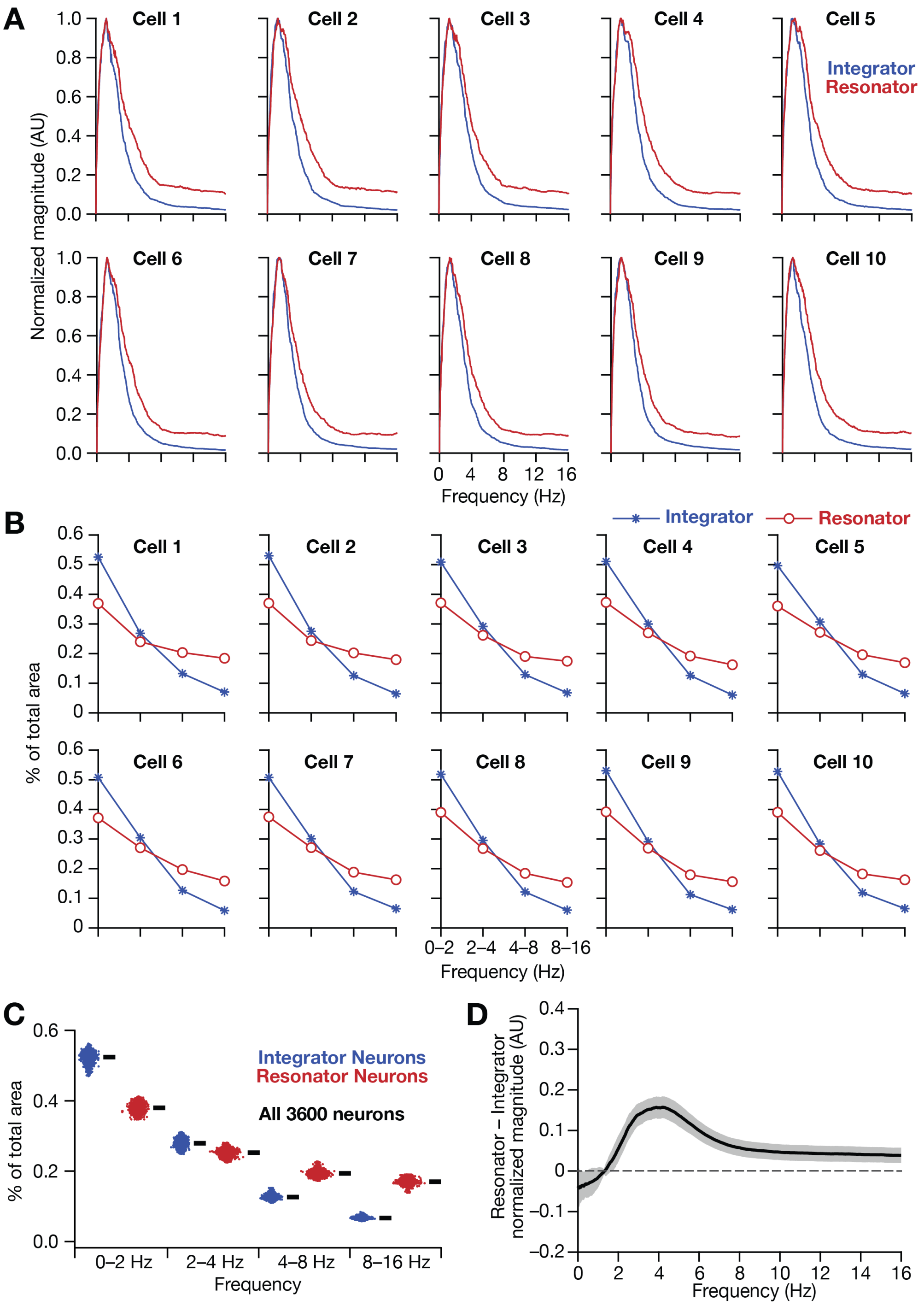

**Figure 13–figure supplement 1. Intrinsically resonating neurons (phenomenological) suppressed low-frequency components and enhanced frequency components around resonance frequency in homogeneous CAN models.** (**A**–**B**) Ten example magnitude spectra (normalized to peak) of grid cell activity (**A**) and the respective percentages of total area covered in each octave of the magnitude spectra (**B**) for homogeneous CAN models with integrator (blue) or resonator (red) neurons. (**C**) Percentage of total area covered in each octave of the magnitude spectra for homogeneous CAN models with integrator (blue) or resonator (red) neurons ($n$ = 3600). Thick black lines represent respective median values. (**D**) Difference between the normalised magnitude spectra of neural temporal activity patterns for integrator and resonator neurons in a homogeneous network.

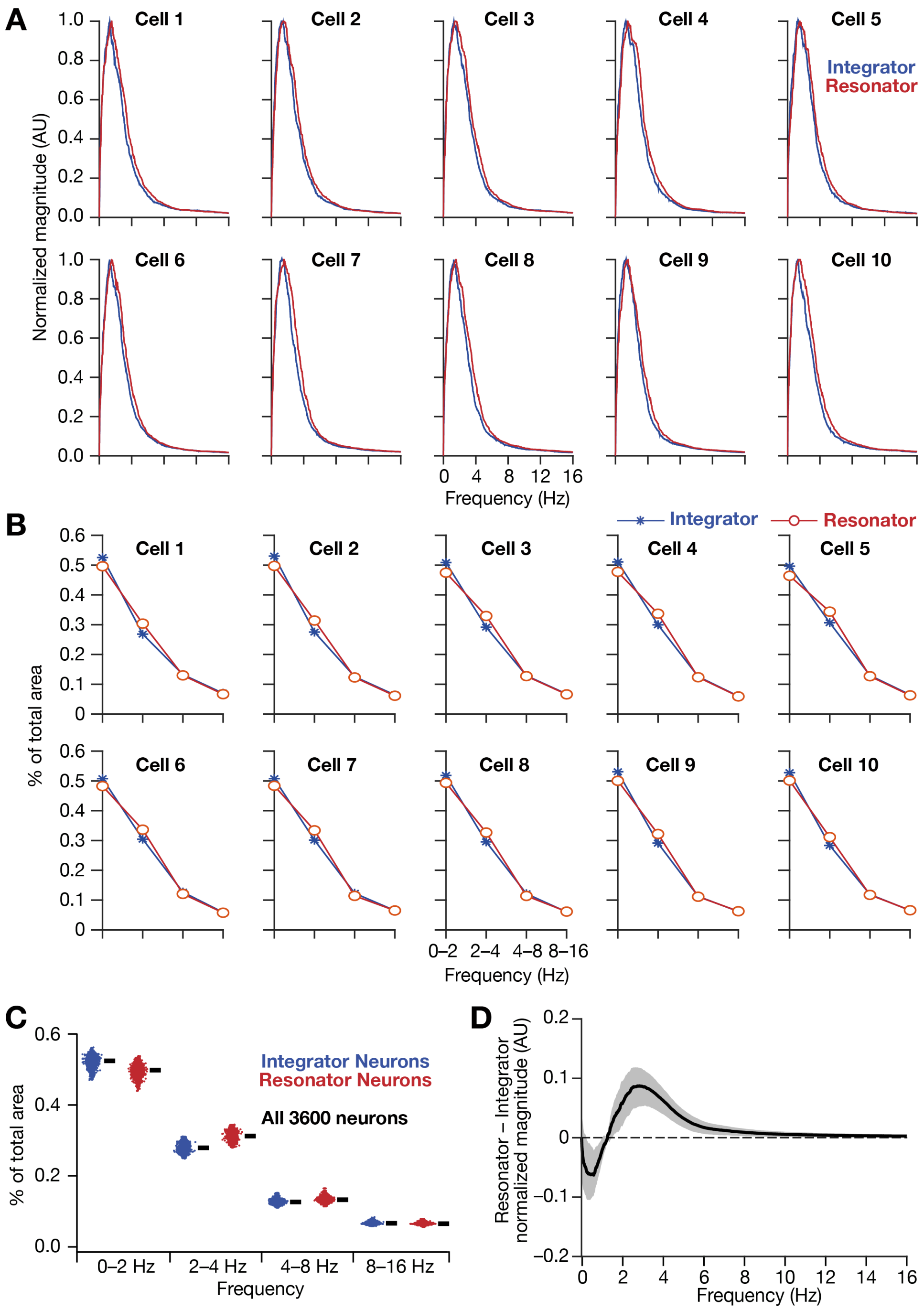

**Figure 13–figure supplement 2. Intrinsically resonating neurons (mechanistic) suppressed low-frequency components and enhanced frequency components around resonance frequency in homogeneous CAN models.** (**A**–**B**) Ten example magnitude spectra (normalized to peak) of grid cell activity (**A**) and the respective percentages of total area covered in each octave of the magnitude spectra (**B**) for homogeneous CAN models with integrator (blue) or resonator (red) neurons. (**C**) Percentage of total area covered in each octave of the magnitude spectra for homogeneous CAN models with integrator (blue) or resonator (red) neurons ($n$ = 3600). Thick black lines represent respective median values. (**D**) Difference between the normalised magnitude spectra of neural temporal activity patterns for integrator and resonator neurons in a homogeneous network. Note that the mechanistic resonator does not introduce spurious high-frequency power as the phenomenological resonator (Figure 13–figure supplement 1). In terms of frequency components of neural activity, the overall impact of the mechanistic resonator on *homogeneous* CAN networks is minimal compared to that of the phenomenological resonator.

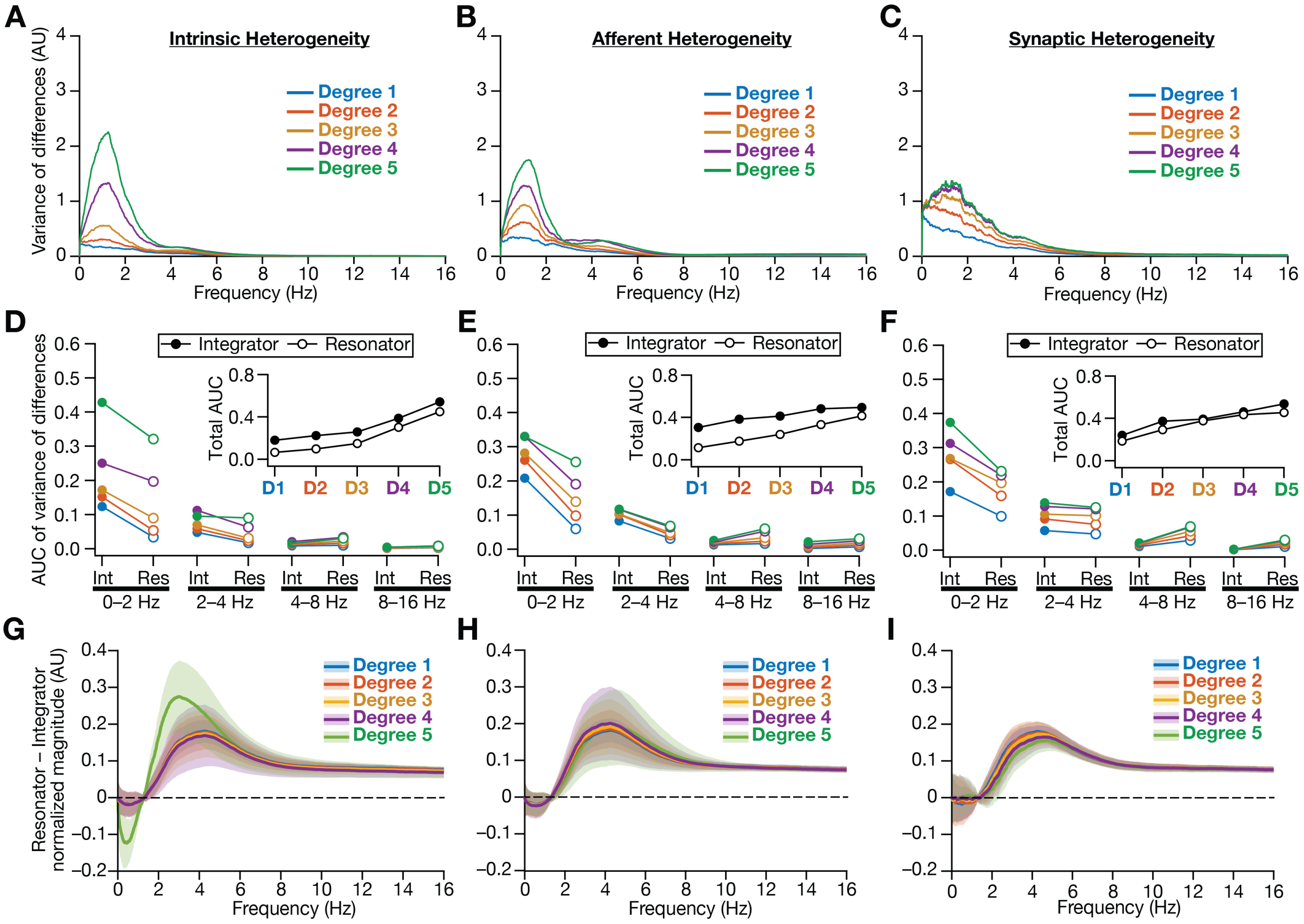

**Figure 13–figure supplement 3. Intrinsically resonating neurons (phenomenological) suppressed heterogeneity-induced variability in low-frequency perturbations caused by different forms of biological heterogeneities.** (**A**–**C**) Normalized variance of the differences between the magnitude spectra of neurons in homogeneous *vs.* heterogeneous networks, across different forms and degrees of heterogeneities, plotted as a function of frequency. (**D**–**F**) Area under the curve (AUC) of the normalized variance plots shown in Figure. 4 (for integrators) and panels A–B (for resonators) showing the variance to be lower in resonator networks compared to integrator networks. Panels depict outcomes for networks with different degrees of intrinsic (**D**), afferent (**E**), and synaptic (**F**) heterogeneities. The respective insets show the total AUC across all frequencies for the integrator *vs.* the resonator networks. (**G**–**I**) Difference between the normalised magnitude spectra of neural temporal activity patterns for integrator and resonator neurons. Panels depict outcomes for networks with different degrees of intrinsic (**G**), afferent (**H**), and synaptic (**I**) heterogeneities. Solid lines depict the mean and shaded area depicts the standard deviations, across all 3600 neurons.

**Figure 2–table supplement 1. Forms and degrees of heterogeneities introduced in the CAN model of grid cell activity.**

| Degree of Heterogeneity | Intrinsic Heterogeneity ($\tau$) | | Afferent Heterogeneity ($\alpha$) | | Synaptic Heterogeneity ($\Delta W_{ij}$) | |
|---|---|---|---|---|---|---|
| | Lower Bound | Upper Bound | Lower Bound | Upper Bound | Lower Bound | Upper Bound |
| 1 | 8 | 12 | 35 | 55 | 0 | 300 |
| 2 | 6 | 14 | 25 | 65 | 0 | 600 |
| 3 | 4 | 16 | 15 | 75 | 0 | 900 |
| 4 | 2 | 18 | 5 | 85 | 0 | 1200 |
| 5 | 1 | 20 | 0 | 100 | 0 | 1500 |
